# Immune Biomarkers of Islet Transplant Rejection Revealed by Synthetic Immunological Niche

**DOI:** 10.64898/2026.05.14.725252

**Authors:** Jyotirmoy Roy, Abdalmonam Jadou Nejma, Mohammad Tarique, Amod Talekar, Shengli Wu, Brianna Ha, Yifei Jiang, Esma S. Yolcu, Lonnie D. Shea

**Affiliations:** Department of Biomedical Engineering, University of Michigan; Ann Arbor, USA; Department of Pediatrics, Department of Molecular Microbiology and Immunology, NextGen Precision Health Institute, Ellis Fischel Cancer Center, University of Missouri; Columbia, USA; Department of Chemical Engineering, University of Michigan; Ann Arbor, USA

## Abstract

Islet transplantation can restore glycemic control in type 1 diabetes, yet the heterogeneity of patient immune responses and transplant outcomes motivates the need for technologies to monitor the graft. Since transplanted islets are not readily accessible for biopsy due to their diffuse engraftment within the liver, clinical monitoring relies on measurements such as islet mass, blood glucose, and C-peptide levels, which are lagging indicators that change only after substantial graft injury. Here, we developed a minimally invasive synthetic immunological niche (IN) that captures graft-associated immune responses through serial subcutaneous biopsy. We evaluated the IN across murine syngeneic, allogeneic, and autoimmune islet transplant models, including CD40/CD154 costimulatory blockade with anti-CD40L. In syngeneic versus allogeneic recipients, IN identified immune populations and transcriptomic signatures that mirrored the graft and distinguished healthy from rejecting grafts. In anti-CD40L treated allografts, IN revealed innate macrophage- and dendritic cell-associated programs linked to graft acceptance versus rejection, whereas IN from untreated allografts showed stronger adaptive immune signatures. Longitudinal IN profiling further detected progressive inflammatory activation in accepted allografts, indicating persistent subclinical risk. Finally, in an autoimmune allograft model treated with anti-CD40L plus rapamycin, IN identified a 13-gene signature that separated early from late rejection trajectories and distinguished autoimmune-from alloimmune-associated rejection programs. Overall, these findings establish IN as a surrogate tissue for minimally invasive monitoring of islet graft and early detection of rejection-associated immune dysregulation.

**One Sentence Summary:** An engineered immunological niche captures distinct immune signatures of allo- and auto-mediated islet transplant rejection

## INTRODUCTION

Islet transplantation offers a transformative treatment option for individuals with brittle autoimmune Type 1 diabetes (T1D), characterized by marked glycemic variability and recurrent episodes of severe, life-threatening hypoglycemia (*1–3*). Islet transplants restore endogenous insulin secretion, thereby improving glycemic stability, and reducing the risk of severe hypoglycemia (*4–8*). Most clinical islet transplants are heavily dependent on some form of immunosuppression to control rejection and recurrent autoimmunity. However, clinical outcomes are still highly heterogeneous, with only 37.5% achieving insulin independence at 1 year, 57-77% maintaining HbA1c <7% long-term, and 69-84% retaining graft function during follow-up (*9–11*). The variable response is primarily because durable graft function remains limited by heterogeneity in immune-mediated loss of transplanted islets (*12*). Unlike solid-organ transplantation such as kidney, liver, and heart, where biopsies, imaging, and better defined circulating biomarkers can aid clinical surveillance, monitoring of islet graft health remains uniquely challenging (*13–16*). Transplanted islets are dispersed, inaccessible to routine biopsy, and have a relatively low tissue volume (*17*). Changes in blood glucose or C-peptide often become evident only after substantial graft injury has already occurred, leaving a narrow or absent window for timely therapeutic intervention and graft salvage (*18, 19*). The delayed clinical detection of rejection and reliance on late functional readouts underscore the need for early biomarkers of immune dysregulation associated with islet graft rejection.

This challenge of monitoring tissue-associated immune dysregulation during disease pathogenesis has been previously addressed with a microporous polycaprolactone scaffold that forms a synthetic immunological niche (IN) following subcutaneous implantation (*20, 21*). The IN recruits host immune and stromal cells, through foreign body response, to generate a vascularized, accessible microenvironment that mirrors systemic and tissue-level immune states, while serving as a non-vital tissue for serial biopsy. Prior studies demonstrated that IN can capture dynamic immune programs across diverse pathological settings, especially autoimmune diseases and organ transplantation (*20, 22, 23*). In T1D, longitudinal profiling of IN in non-obese diabetic (NOD) mice enabled identification of early myeloid-associated immune dysregulation followed by lymphoid dysregulation at later stages of disease progression, revealing inflammatory signatures that precede overt hyperglycemia (*24*). These findings further demonstrated the ability of IN to capture critical tissue-associated immune transcriptomic changes and, in turn, to monitor efficacy of immunotherapies. The utility of IN was further supported in models of heart and skin transplantation, where it detected immune signatures associated with acute cellular allograft rejection (*22*). These signatures captured immune activation that preceded overt graft injury and rejection. Collectively, these studies highlight the advantages of IN biopsy over peripheral blood monitoring, as circulating immune cells are often relatively naive and resting, whereas tissue-resident and tissue-recruited immune cells are more activated, functionally conditioned, and therefore more reflective of ongoing disease processes (*25, 26*).

In this report, we investigated the utility of IN for monitoring islet auto-allograft rejection under anti-CD40L immunotherapy, and the ability to distinguish autoimmune from alloimmune components of rejection. Tegoprubart, an anti-CD40L monoclonal antibody recently evaluated in clinical studies, has demonstrated promising outcomes in islet transplantation using CD40/CD154 pathway blockade, supporting its translational relevance as an immunomodulatory strategy (*27*). The utility of IN was first assessed in syngeneic and allogeneic islet transplant models, with INs implanted subcutaneously at time of transplantation. IN captured innate and adaptive immune cell populations relevant to islet rejection and identified a gene signature distinguishing syngeneic from allogeneic recipients that paralleled immune changes within graft, supporting its use as a surrogate tissue for graft monitoring. We next profiled IN biopsies from accepted and rejected allografts under anti-CD40L treatment. Transcriptomic and flow cytometric analyses revealed macrophage- and dendritic cell-associated populations linked to graft acceptance relative to rejection. INs further demonstrated distinct rejection mechanisms, with anti-CD40L treated allografts exhibiting predominantly innate immune-driven rejection programs, in contrast to untreated allografts, which showed stronger adaptive immune signatures. INs also captured increasing immune inflammatory signals over time in accepted allografts relative to syngeneic controls, suggesting ongoing risk for later graft failure. Finally, in an autoimmune-allograft model treated with anti-CD40L and rapamycin, IN identified a 13-gene signature that distinguished early from late rejection trajectories. These analyses further suggested that adaptive immune programs were associated with autoimmunity, whereas innate immune programs were more strongly linked to alloimmune responses under anti-CD40L. Together, these findings demonstrate that the minimally invasive IN platform can capture otherwise inaccessible graft-associated immune dysregulation, potentially enabling earlier intervention and improved long-term preservation of islet graft function in clinical settings.

## RESULTS

### IN captures allogeneic vs syngeneic immune differences mirrored in islet grafts

We applied IN in an islet transplant model to capture immune differences between rejecting allogeneic and stable syngeneic grafts. Diabetes was induced in C57BL/6 mice using streptozotocin, followed by transplantation of either allogeneic islets (BALB/c to C57BL/6) or syngeneic islets (C57BL/6 to C57BL/6). INs were implanted subcutaneously at time of transplant and serially biopsied with replacement from day 7-70 or onset of hyperglycemia (**Fig. 1a**). Allogeneic recipients without immunosuppression showed an initial reduction in blood glucose level following transplantation; however, all mice subsequently rejected the engrafted islet cells and returned to overt hyperglycemia (≥250 mg/dL) by day 14 (**Fig. 1b, c, Sup. Table 1**). In contrast, syngeneic recipients maintained normoglycemia through day 70, indicating durable graft function (**Fig. 1b, c**). We next examined IN immune composition in the two groups at day 14 using spectral flow cytometry (**Sup. Fig. 1, Sup. Table 2)**. Relative to syngeneic controls, allogeneic INs contained a greater proportion of total immune cells, consistent with heightened immune recruitment during graft rejection (**Fig. 1d**). In the immune cell compartment, reduced neutrophil frequencies, and a trend toward increased macrophage abundance in allogeneic group was observed (**Fig. 1e, f**). Dendritic cell proportions were similar among both groups (**Fig. 1g)**. Although total T cell proportions were comparable, allogeneic INs showed enrichment of CD4 T cells within the T cell compartment, whereas CD8 T cell frequencies were unchanged (**Fig. 1h-j**), supporting a stronger helper/effector adaptive response at IN during rejection. Syngeneic INs had an increase in CD43+CD8+ T cells, while CD43+CD4+ T cells were unchanged (**Fig. 1k, l**). Since allogeneic grafts had developed hyperglycemia by day 14, this difference may reflect contraction or redistribution of activated CD8 T cells after peak graft injury. Syngeneic INs also showed higher proportions of total B cells and CD43+ B cells, consistent with recent studies highlighting a potential regulatory role of these B cell phenotypes (*28*) (**Fig. 1m-n**). Most of these lymphoid subset populations were a small fraction of total immune cells. Notably, comparison of IN and liver graft-site immune composition at respective end timepoints showed that relative allogeneic-versus-syngeneic trends were largely preserved across both tissues (**Sup. Fig. 2, Sup. Table 3**).

**Figure 1.**
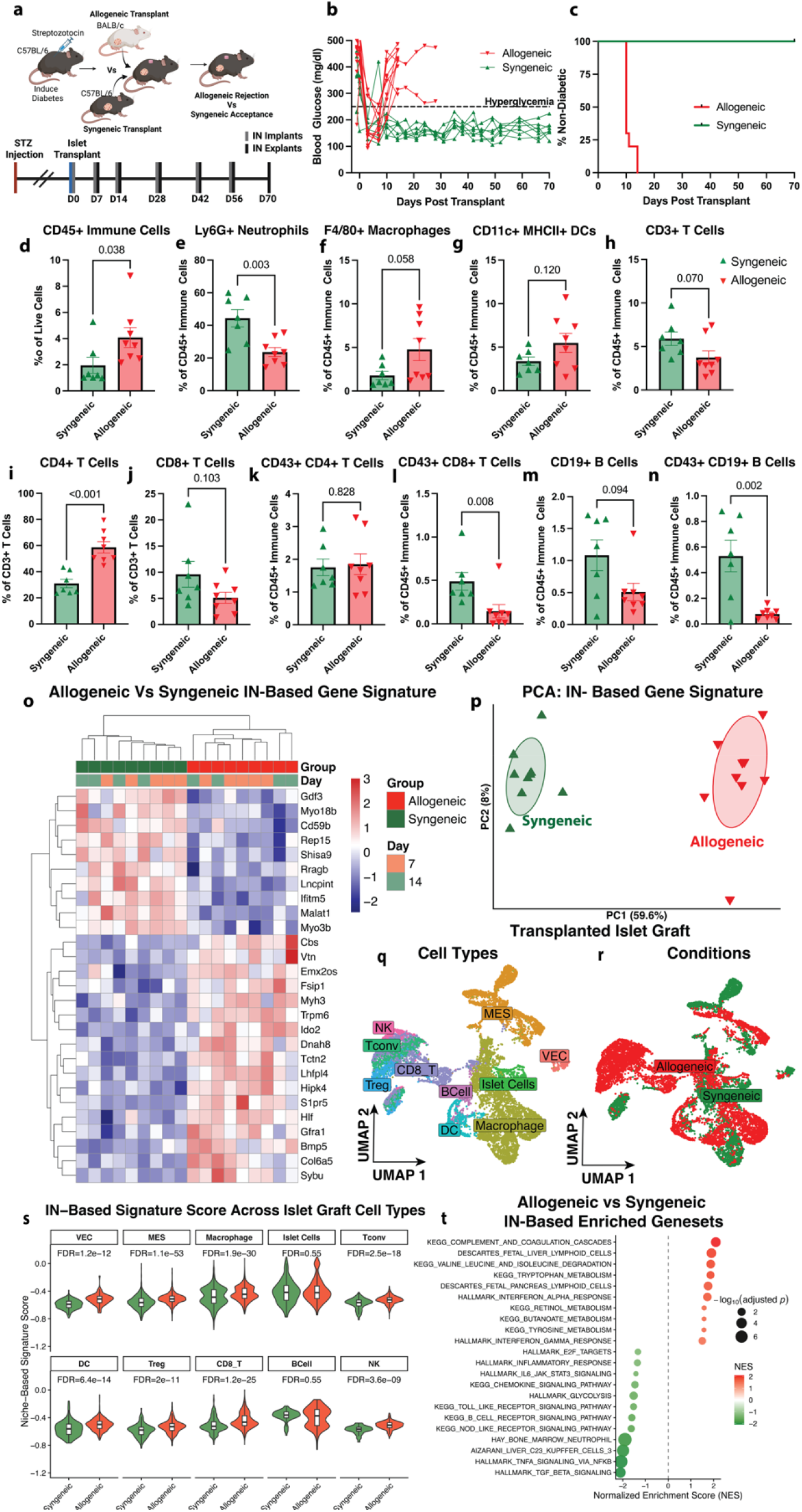
IN profiling distinguishes allogeneic and syngeneic islet transplants and reflects graft-associated immune signature. **(a)** Experimental design: Streptozotocin-induced diabetic C57BL/6J mice received allogeneic (BALB/c) or syngeneic (C57BL/6J) islet transplants with concurrent subcutaneous IN implantation. INs were longitudinally biopsied with replacement until day 70 post-transplant or onset of hyperglycemia. **(b)** Blood glucose trajectories, and **(c)** Diabetes-free survival demonstrating rejection of allogeneic grafts (n=10) by day 14, whereas syngeneic grafts (n=7) remained normoglycemic through day 70. Proportions of **(d)** immune cells, **(e)** neutrophils, **(f)** macrophages, **(g)** dendritic cells, **(h)** T cells, **(i)** CD4+ T cells, **(j)** CD8+ T cells, **(k)** CD43+ CD4 T cells, **(l)** CD43+ CD8+ T cells, **(m)** B cells, and **(n)** CD43+ B cells in allogeneic (n=8) and syngeneic (n=7) INs at day 14. Data shown as mean ± SEM. Statistical comparisons used unpaired t-tests or Mann-Whitney tests based on normality. **(o)** Heatmap and **(p)** PCA showing Elastic Net-derived IN transcriptomic signature separating allogeneic and syngeneic samples (n=4-5/group/timepoint) across days 7 and 14. **(q)** Cell type and **(r)** condition distribution from day 7 islet graft single-cell RNA-seq. **(s)** IN-derived signature scores across graft cell types. **(t)** GSEA of IN highlighting selected pathways enriched between allogeneic and syngeneic conditions.

We next examined in-depth phenotypic differences within IN using bulk RNA sequencing of IN biopsies collected at days 7 and 14 post-transplant. After preprocessing and filtering, feature selection using an elastic net model (*α*= 0.5) identified a 27-gene signature that robustly distinguished allogeneic from syngeneic recipients (**Fig. 1o**). Genes enriched in syngeneic group included *Cd59b*, a complement regulatory molecule that protects against complement-mediated injury (*29*); *Malat1*, a gene shown to regulate inflammation in autoimmune diseases (*30*); and *Lncpint*, which is associated with immune homeostasis and inflammation control (*31*). Together, these features suggest that syngeneic grafts are associated with a more tissue-protective immune state. In contrast, genes elevated in allogeneic INs were associated with more adaptive immune and pro-inflammatory response, like *S1pr5*, linked to T cell trafficking, *Hlf*, linked to promoting pro-inflammatory activity in CD4+ T Cells, and *Ido2*, implicated in pro-inflammatory mediator of B cell responses (*32–34*). Principal component analysis based on this gene signature showed clear separation of the two transplant settings across both time points (**Fig. 1p)**, demonstrating that IN captures transcriptomic states associated with allogeneic relative to syngeneic groups. The changes in IN-based immune transcriptome was reflective of the biology within the transplanted islet graft. We mapped the IN-derived transcriptomic signature onto a published single-cell RNA sequencing dataset of syngeneic and allogeneic islet grafts collected at day 7 post-transplant (*35*). Analysis of the graft dataset identified diverse immune and stromal populations, including macrophages, dendritic cells, T cell subsets, islet cells, endothelial cells, and mesenchymal cells from both the allogeneic and syngeneic transplant settings (**Fig. 1q, r**). We then generated an IN-based signature score across graft cell populations by calculating the difference between the mean expression of allogeneic-enriched and syngeneic-enriched genes within the IN signature. Strikingly, the IN-based signature score was significantly elevated across majority of cell types in allogeneic relative to syngeneic condition (**Fig. 1s**). This suggested that transcriptomics state captured by distant subcutaneous IN closely mirrored those within transplanted islet graft, supporting that IN can function as a surrogate tissue for islet graft.

Pathway-level analysis of IN by gene set enrichment indicated enrichment of lymphoid and IFN-γ inflammatory programs in allogeneic condition (**Fig. 1t, Sup. Table 4**). In contrast, syngeneic group was enriched for neutrophil, TGF-β, TNF-α/IL-6, and NOD-like receptor signaling pathways, suggesting a mixed inflammatory and tissue-associated immune response. Similar pathway trends were observed at graft site as well (**Sup Fig. 3**). Together, these data indicate that IN captures not only cellular and gene-level differences, but also broader immune program states associated with allogeneic rejection versus syngeneic engraftment.

### IN identifies innate immune signatures of allograft acceptance vs rejection under anti-CD40L

We next investigated immune differences within IN in context of immunosuppression by comparing accepted and rejected islet allografts under anti-CD40L immunotherapy. Streptozotocin-induced diabetic C57BL/6 mice received allogeneic islets (BALB/c to C57BL/6) together with three 500*µ*g doses of anti-CD40L on days 0, 7, and 14. INs were implanted subcutaneously at time of transplantation and serially biopsied with replacement from day 7 until day 70 or onset of hyperglycemia (**Fig. 2a**). Most anti-CD40L treated recipients (14/17) showed reduced blood glucose after transplantation, accepted the graft, and maintained normoglycemia through day 70 (**Fig. 2b, c, Sup. Table 1**). A few mice (n=3) which were observed for long term showed increase in blood glucose and graft rejection only after 100 days post-transplant (**Fig. 2c**). In contrast, anti-CD40L treated rejecting allograft recipients exhibited an initial improvement followed by recurrent hyperglycemia by day 14 (**Fig. 2c**). We analyzed IN immune composition from anti-CD40L accepted and rejected allotransplant groups at day 14 using spectral flow cytometry (**Sup. Table 2**). Relative to accepted group, INs from rejected group had a higher proportion of total immune cells, consistent with higher immune recruitment related to rejection (**Fig. 2d**). This increase was accompanied by increased neutrophil proportion, suggesting a more inflammatory niche environment (**Fig. 2e**). In contrast, INs from accepted allografts were enriched for macrophages, particularly CX3CR1+ and CD80+ macrophages, suggesting that graft acceptance is associated with specialized myeloid programs involving immune regulation, tissue surveillance, and antigen-presenting function (**Fig. 2f-h)**. A similar trend was observed for total dendritic cells and CX3CR1+ dendritic cells, which were also higher in INs in accepted group (**Fig. 2i, j)**. No significant differences were observed in total T cells or in proportions of CD4+ and CD8+ T-cell between the groups (**Fig. 2k-m**). Accepted allografts also showed higher frequencies of total B cells and CD43+ B cells, indicating additional adaptive immune features associated with durable graft survival (**Fig. 2n, o**). Interestingly, the IN continued to reflect graft-associated immune changes, as both the IN and liver graft site exhibited similar compositional shifts from untreated to anti-CD40L treated allogeneic conditions (**Sup. Fig. 4, Sup. Table 3**).

**Figure 2.**
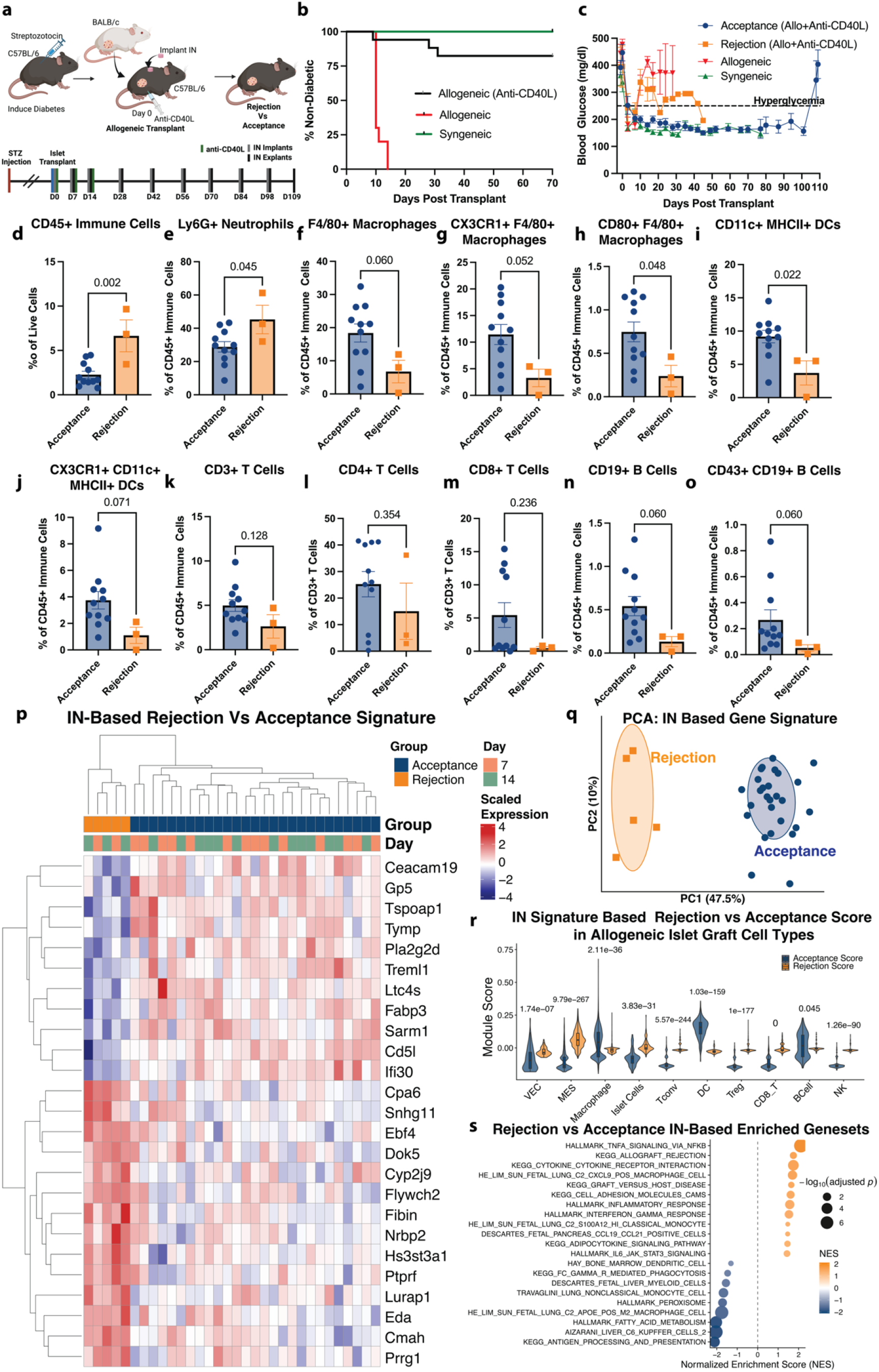
IN identifies innate immune programs distinguishing with allograft acceptance and rejection under anti-CD40L therapy. **(a)** Experimental design: Streptozotocin-induced diabetic C57BL/6J mice received allogeneic (BALB/c) islet transplants with concurrent subcutaneous IN implantation and anti-CD40L treatment. INs were longitudinally biopsied with replacement until day 70 post-transplant or onset of hyperglycemia. A subset of mice (n=3) was monitored through day 109 post-transplant. **(b)** Diabetes-free survival in allogeneic (n=10), syngeneic (n=7), and allogeneic + anti-CD40L (n=17) groups. **(c)** Blood glucose trajectories of accepted (n=14) and rejected (n=3) anti-CD40L-treated allografts. Proportions of **(d)** immune cells, **(e)** neutrophils, **(f)** macrophages, **(g)** CX3CR1+ macrophages, **(h)** CD80+ macrophages, **(i)** dendritic cells, **(j)** CX3CR1+ dendritic cells, **(k)** T cells. **(l)** CD4+ T cells, **(m)** CD8+ T cells, **(n)** B cells, and **(o)** CD43+ B cells in accepted (n=11) and rejected (n=3) INs at day 14, shown as mean ± SEM. Statistical comparisons used unpaired t-tests or Mann-Whitney tests based on normality. **(p)** Heatmap and **(q)** PCA showing Elastic Net-derived transcriptomic signature separating rejection (n=2-3/timepoint) and acceptance (n=13-14/timepoint) samples across days 7 and 14. **(r)** IN-signature based acceptance and rejection scores across allograft cell types. **(s)** GSEA of IN highlighting selected pathways enriched between accepted and rejected conditions.

We identified that IN-based immune transcriptomic changes are reflective of biological differences between accepted and rejected allograft under anti-CD40L immunotherapy. Bulk RNA sequencing was performed on IN biopsies collected at days 7 and 14. Following preprocessing and filtering, elastic net feature selection (*α*= 0.3) identified a 25-gene signature that distinguished graft acceptance from rejection (**Fig. 2p**). Genes enriched in the accepted group included *Pla2g2d*, an immunoregulatory lipid mediator related to M2-like macrophages (*36*); *Treml1*, associated with myeloid related inflammation (*37*); *Ltc4s*, involved in leukocyte inflammatory mediator pathways (*38*); *Cd5l*, associated with macrophage for immune regulation (*39*); and *Ifi30*, an antigen-processing gene promoting M2-like macrophage polarization (*40*). Collectively, these genes suggest that graft acceptance is characterized by a controlled myeloid state that integrates immune surveillance, antigen handling, and inflammation regulatory programs. In contrast, rejecting grafts showed higher expression of genes including *Lurap1*, linked to activation of canonical NF-kappa-B pathway and pro-inflammatory cytokine production (*41*); and *Ebf4*, a regulator of cytotoxic functions in immune cells (*42*). Principal component analysis based on this signature demonstrated clear separation of accepted and rejected samples across both time points (**Fig. 2o**), highlighting the ability of IN to resolve early transcriptomic states linked to divergent allo-transplant outcomes under anti-CD40L therapy.

IN-based acceptance versus rejection signature under anti-CD40L was also reflected within islet allograft microenvironment, particularly in myeloid compartments. We mapped IN-derived signature onto single-cell RNA sequencing data from allogeneic islet grafts collected at day 7 post-transplant (*35*). Acceptance and rejection scores were calculated across all allograft cell populations using module scores derived from IN signature genes enriched in each state. Notably, macrophages and dendritic cells exhibited higher acceptance scores than rejection scores (**Fig. 2r**), indicating that myeloid programs identified in IN are recapitulated at graft site. This finding was consistent with our earlier flow cytometry data showing enrichment of anti-inflammatory CX3CR1+ macrophages and dendritic cells in accepted grafts. Functional relevance of these myeloid states was further supported by gene set enrichment analysis of IN in anti-CD40L treated accepted vs rejected allografts groups (**Sup. Table 4)**. Rejected grafts showed relatively higher enrichment of inflammatory pathways, including TNF-*α*and interferon-*γ*signaling, together with signatures of classical monocytes and CXCL9-associated inflammatory macrophages (**Fig. 2s**). In contrast, accepted grafts were enriched for nonclassical monocytes, M2-like macrophages, and antigen-presentation programs, supporting a model in which graft survival is associated with regulated and anti-inflammatory myeloid immunity.

### Allograft rejection under anti-CD40L is innate immune driven vs T cell driven rejection in control

We next investigated the different immune mechanisms driving allograft rejection with and without anti-CD40L immunotherapy. INs from anti-CD40L treated rejected allografts were compared with INs from untreated rejected allografts in C57BL/6 model (**Fig. 3a**). Flow cytometry analysis at day 14 revealed that IN from anti-CD40L treated rejected allografts mice contained higher frequencies of neutrophils, with trends toward increased CD80+ dendritic cells, and CCR2+ dendritic cells than untreated controls (**Fig. 3b-d**), indicating a rejection program dominated by innate inflammatory recruitment and activated antigen-presenting cells. In contrast, INs from the untreated control group had higher proportions of B cells, CD4+ T cells, and CD8+ T cells (**Fig. 3e-g**), consistent with a more lymphocyte-driven rejection response when anti-CD40L is absent. Together, these findings suggest that anti-CD40L reshapes alloimmune rejection by shifting dominant biology from adaptive T cell responses toward innate immune mechanisms.

**Figure 3.**
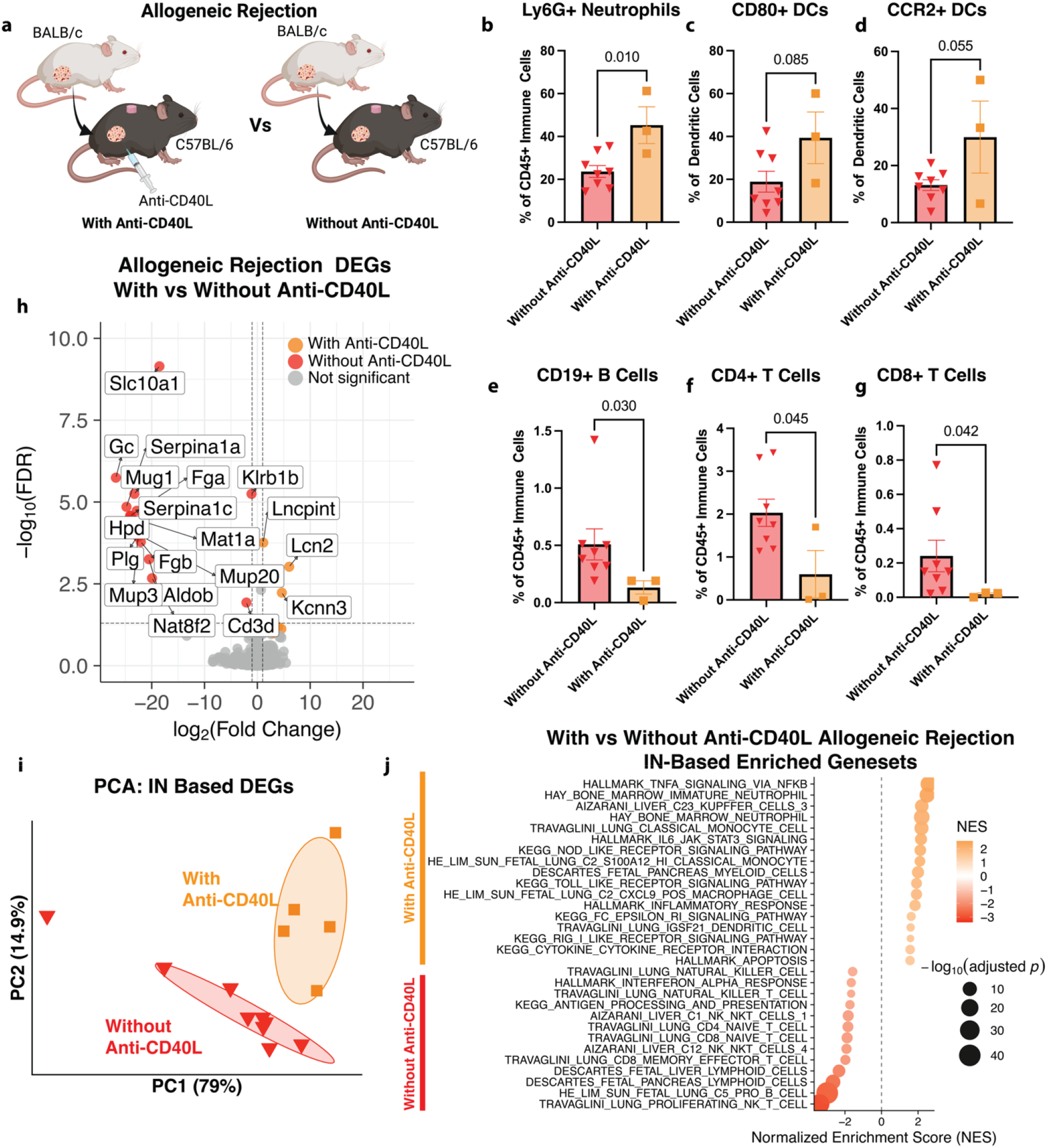
Allograft rejection under anti-CD40L is innate immune-driven vs T cell-driven rejection in control. **(a)** Experimental design: INs from anti-CD40L treated rejected allografts (days 7 and 14) were compared with untreated allogeneic controls. **(b-g)** Spectral flow cytometry of INs at Day 14 reveals distinct immune composition between anti-CD40L treated (n=3) and untreated (n=8) conditions, including **(b)** neutrophils, **(c)** CD80+ dendritic cells, **(d)** CCR2+ dendritic cells, **(e)** B cells, (**f)** CD4+ T cells, and **(g)** CD8+ T cells. Statistical comparisons were performed using unpaired t-tests (**b, c, d, f**) or Mann-Whitney tests (**e, g**) based on Shapiro-Wilk normality testing, with data shown as mean ± SEM. **(h)** Volcano plot highlighting differentially expressed innate and adaptive immune genes between anti-CD40L treated rejected allografts (n=2-3 per timepoints) and untreated controls (n=4-5 per timepoints) at Day 7 and 14 combined. Axes truncated for visualization. **(i)** PCA of differentially expressed genes showing separation between groups (70% confidence ellipses). **(j)** Selected differentially enriched pathways between anti-CD40L treated rejection and control allogeneic conditions from Gene set enrichment analysis.

IN transcriptomic profiling further confirmed that rejection under anti-CD40L was more innate immune driven, whereas rejection in untreated control allografts was dominated by adaptive immunity. Differential gene expression analysis of IN biopsies collected at days 7 and 14 identified genes associated with distinct rejection programs (|log2FC| ≥ 1, padj ≤ 0.1, **Fig. 3h, Sup. Table 5**). Untreated control allografts showed higher expression of adaptive immune genes such as *Cd3d* and *Klrb1b*, consistent with T- and lymphocyte-mediated rejection. In contrast, INs from anti-CD40L treated rejected allografts were enriched for innate immune genes including *Lcn2* and *Lncpint*, supporting persistence of a myeloid-related inflammatory state despite costimulatory blockade. PCA using the differentially expressed genes showed distinct clustering of anti-CD40L treated rejected allografts from control untreated allografts (**Fig. 3i**).

Gene set enrichment analysis reinforced the differential rejection programs (**Sup. Table 4)**. Anti-CD40L treated rejected allografts showed higher enrichment of neutrophil, classical monocyte, and CXCL9-associated inflammatory macrophage programs, together with TNF-*α* inflammatory pathways (**Fig. 3j**). In contrast, untreated control allografts were enriched for lymphoid programs including CD4+ T cells, CD8+ T cells, B cells, and NKT cell gene sets. Collectively, these results demonstrate that anti-CD40L alters dominant biology of rejection from adaptive lymphocyte responses toward innate inflammatory mechanisms, highlighting the ability of IN to resolve mechanistically distinct pathways of graft failure.

### Allograft acceptance under anti-CD40L has distinct immune signature over time from syngeneic

We investigated differences in immune state of graft acceptance under anti-CD40L from control syngeneic graft tolerance, to identify residual susceptibility to future rejection. Longitudinal IN biopsies from anti-CD40L treated accepted allografts were therefore compared with INs from syngeneic controls in C57BL/6 model (**Fig. 4a**). Flow cytometry of IN was performed at days 14, 42, and 70 to define how immune states evolved over time in the two groups. INs from accepted allografts showed a progressive increase in total immune cell abundance over time (**Fig. 4b**), suggesting increasing immune engagement despite preserved graft function. Neutrophils declined over time in syngeneic group but remained relatively stable in accepted allografts (**Fig. 4c**), consistent with persistent inflammation profile in the allogeneic acceptance group. Dynamic changes were also observed in other myeloid compartments. In the syngeneic group, the proportion of total macrophages increased over time, including anti-inflammatory CX3CR1+ macrophages. In contrast, accepted allografts had a decline in total macrophages and CX3CR1+ macrophages over time (**Fig. 4d, e**). A similar pattern was observed for dendritic cells. INs from syngeneic controls demonstrated a gradual increase in dendritic cell frequencies over time, including an increase in anti-inflammatory CX3CR1+ dendritic cells. In contrast, INs from accepted allografts maintained relatively stable frequencies of these populations at lower numbers **(Fig. 4f, g)**. Although CD4+ T cells increased similarly in both groups, CD8+ T cells expanded more prominently in accepted allografts than in syngeneic controls (**Fig. 4h, i**), indicating increasing adaptive immune activity over time despite initial allograft acceptance.

**Figure 4.**
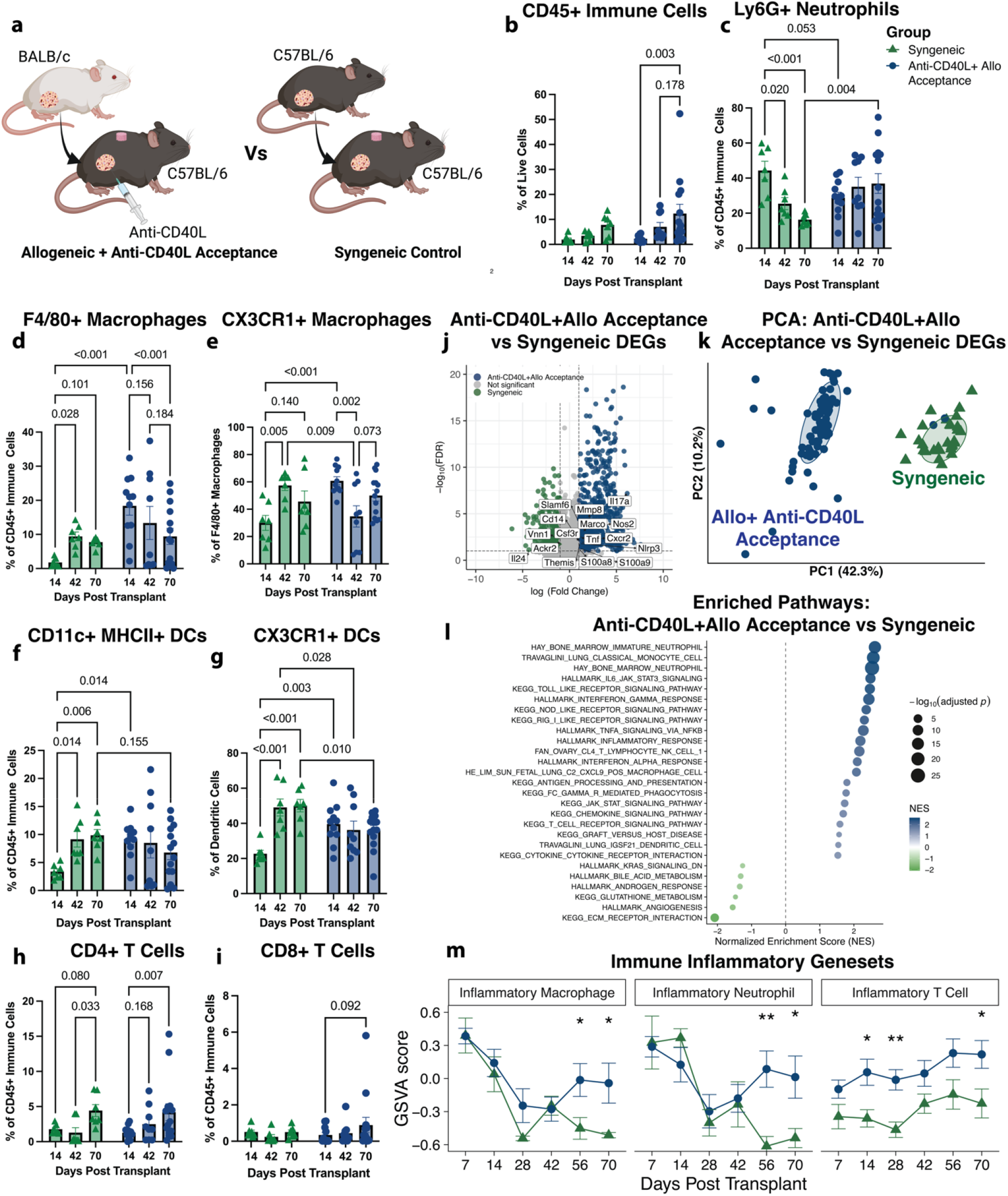
Allograft acceptance under anti-CD40L has distinct immune signatures from syngeneic controls. **(a)** Experimental design: INs from anti-CD40L-treated accepted allografts (days 7, 14, 28, 42, 56, 70) were compared with syngeneic controls. **(b–i)** Longitudinal immune profiling of INs at days 14, 42, and 70 showing temporal immune changes between syngeneic (n=7/timepoint) and anti-CD40L-treated accepted allograft (n=9-14/timepoint) groups, including **(b)** immune cells, **(c)** neutrophils, **(d)** macrophages, **(e)** CX3CR1+ macrophages, **(f)** dendritic cells, **(g)** CX3CR1+ dendritic cells, **(h)** CD4+ T cells, and **(i)** CD8+ T cells. Statistical analysis used mixed-effects models with Tukey’s multiple-comparison test. Data shown as mean ± SEM. **(j)** Volcano plot of differentially expressed innate and adaptive immune genes between anti-CD40L-treated accepted allografts (n=11-14/timepoint) and syngeneic controls (n=4-5/timepoint) across days 7-70. Axes truncated for visualization. **(k)** PCA of differentially expressed genes with 70% confidence ellipses showing separation between groups. **(l)** Selected enriched pathways from GSEA of IN between anti-CD40L-treated accepted and syngeneic conditions. **(m)** Sample-averaged GSVA scores for inflammatory macrophage, neutrophil, and T cell signatures at IN across days 7-70. Welch’s t-test performed. Mean ± SEM shown.

Transcriptomic differences of INs from anti-CD40L treated accepted allografts versus syngeneic controls further demonstrated that accepted allograft group remains immunologically distinct from syngeneic group. Differential gene expression analysis of longitudinal IN biopsies from Day 7 to 70 identified differentially expressed genes between the two groups (|log2FC| ≥ 1, padj ≤ 0.1; **Fig. 3j, Sup. Table 6**). Accepted allografts had higher expression of both innate and adaptive immune genes, including pro-inflammatory cytokine signaling genes such as *Il17a, Tnf*, and *Nos2*, neutrophil recruitment and activation genes such as *Cxcr2, Mmp8*, and *Csf3r*, innate macrophage genes such as *Marco* and *Cd14*, and, T-cell signaling gene like *Themis* (*43–51*). Principal component analysis based on these differentially expressed genes demonstrated clear separation between accepted allografts and syngeneic controls (**Fig. 3k**), indicating persistent immune divergence despite maintained graft function.

Pathway-level analysis at IN supported immune divergence between anti-CD40L treated accepted allografts vs syngeneic controls (**Sup. Table 4)**. INs from accepted allografts were enriched for gene sets related to interferon, IL-6, and TNF-α inflammatory signaling, together with antigen presentation, neutrophil, T-cell receptor, and classical monocyte programs, indicating higher immune inflammatory state (**Fig. 4l**). In contrast syngeneic controls were enriched for angiogenesis, consistent with tissue repair and stable graft integration (**Fig. 4l**). To further examine longitudinal immune trajectories, we quantified gene set variation scores for inflammatory macrophage, neutrophil, and T-cell signatures over time (**Sup. Table 7)**. Inflammatory macrophage and neutrophil programs were initially similar between groups but began to diverge at later time points, particularly from day 56 onward, increasing in accepted allograft group while declining in syngeneic controls (**Fig. 4m**). This pattern suggests that accepted allografts may remain at risk of future rejection rather than progressing toward full immune tolerance. In addition, inflammatory T-cell signatures remained consistently higher in accepted allografts throughout the study, further supporting presence of sustained alloimmune activity despite apparent allograft acceptance (**Fig. 4m**).

### IN captures immune biomarkers of Auto-Allo transplant rejection

We next incorporated the autoimmune component of clinical islet transplantation by performing allogeneic islet transplantation in diabetic autoimmune NOD mice (C57BL/6 to NOD), thereby modeling combined alloimmune and autoimmune response faced by transplanted grafts (**Fig. 5a**). INs were implanted at a pre-diabetic stage. In this more stringent setting, treatment with three doses of anti-CD40L alone was insufficient, and recipients uniformly rejected the graft by day 14 (**Fig. 5b**). In contrast, intensification of immunotherapy with additional rapamycin (2 mg/kg, 15/20 doses) or a fourth anti-CD40L dose improved graft outcomes and significantly delayed rejection (**Fig. 5b**). Blood glucose trajectories in anti-CD40L plus rapamycin groups further revealed two biologically distinct response patterns: an early rejection group that rejected graft and became hyperglycemic by day 33 and a late rejection group that rejected after day 41 (**Fig. 5c, Sup. Table 1**). We profiled IN immune composition at day 14 to detect early immune composition associated with these divergent outcomes. For both groups, INs were predominantly composed of myeloid cells, particularly neutrophils, macrophages, and dendritic cells, with smaller contributions from T and B cells and no significant differences in overall immune cell frequencies between groups (**Fig. 5d-i**). Comparative analysis of IN and liver graft-site immune composition at matched end timepoints showed that early versus late rejection patterns were largely consistent across tissues, further supporting the ability of the IN to capture graft-associated immune response in the autoimmune allograft setting (**Sup. Fig. 5, Sup. Table 3**). These findings also suggested that conventional immune composition alone may not fully explain divergent graft trajectories in auto-allo setting, motivating deeper transcriptomic interrogation of IN.

**Figure 5.**
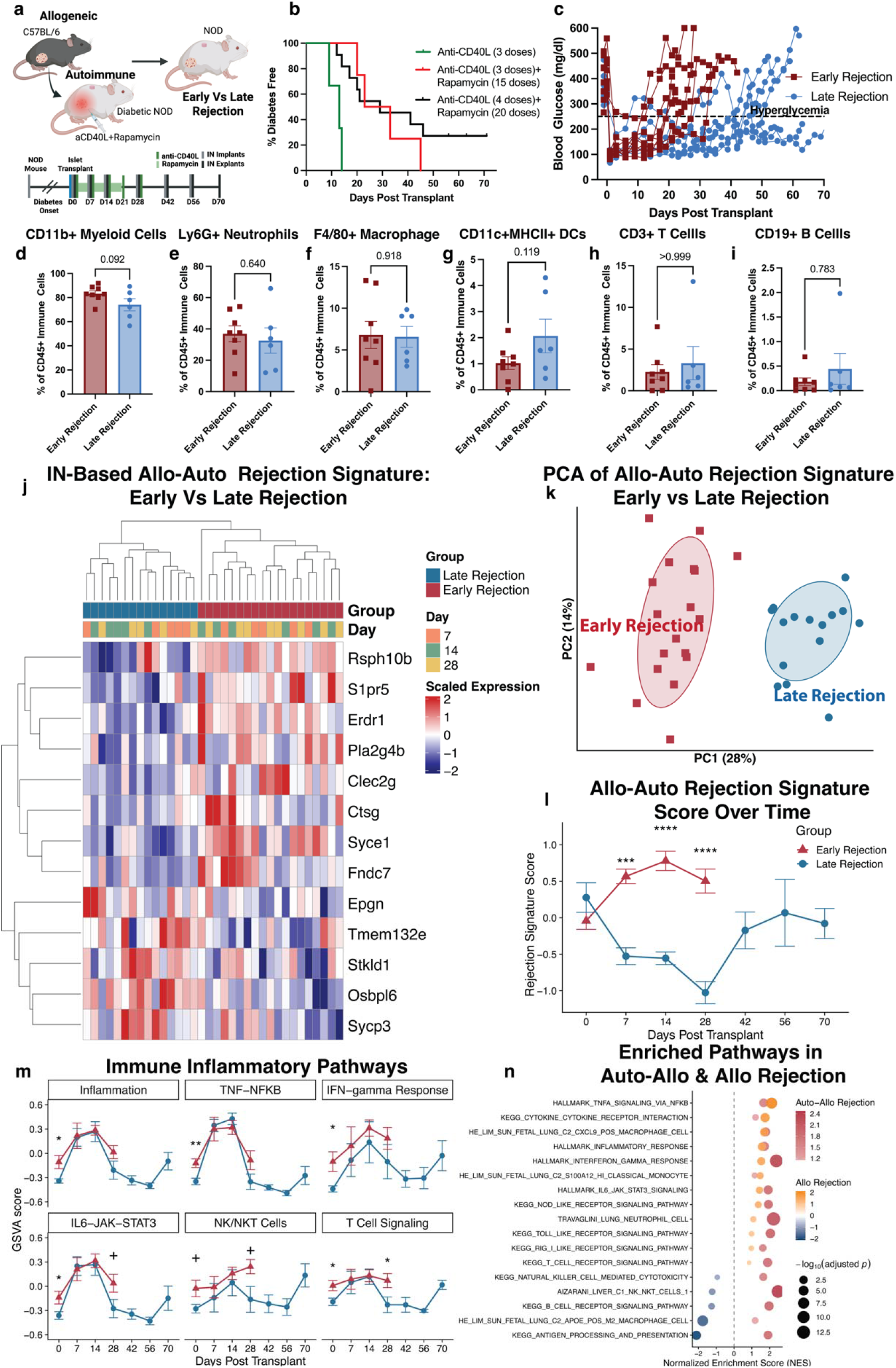
INs capture immune transcriptomic signatures associated with auto-allogeneic rejection. **(a)** Experimental design: Prediabetic NOD mice received subcutaneous IN implantation before diabetes onset, followed by allogeneic islet transplantation (C57BL/6J) after diabetes onset with anti-CD40L and rapamycin treatment. INs were longitudinally biopsied with replacement until day 70 post-transplant or onset of hyperglycemia. **(b)** Diabetes-free survival for anti-CD40L alone (3 doses, n=3), anti-CD40L (3 doses) + rapamycin (15 doses, n=4), and anti-CD40L (4 doses) + rapamycin (20 doses, n=11). **(c)** Blood glucose trajectories identifying early (≤Day 33, n=9) and late (≥Day 41, n=6) rejection. Proportions of **(d)** myeloid cells, **(e)** neutrophils, **(f)** macrophages, **(g)** dendritic cells, **(h)** T cells, **(i)** B cells in IN from early (nado) and late (n=6) rejection groups at day 14. Data shown as mean ± SEM. Statistical comparisons used unpaired t-tests or Mann-Whitney tests based on normality. **(j)** Heatmap and **(k)** PCA showing a 13-gene Elastic Net signature separating early (n=5-8/timepoint) and late (n=5/timepoint) rejection across days 7, 14, and 28. **(l)** Longitudinal rejection scores derived from the 13-gene signature (n=4-8/group/timepoint). **(m)** GSVA inflammatory pathway scores across days 0-70. **(n)** Comparison of selected immune pathways between auto-allogeneic and allogeneic rejection.

Transcriptomic profiling of IN from early- and late-rejection groups identified a distinct auto-allo rejection signature associated with divergent graft outcomes. Following preprocessing and filtering of IN bulk RNA sequencing data from days 7, 14, and 28, elastic net feature selection (α = 0.6) defined a 13-gene signature that separated early rejection from late rejection in the autoimmune-allogeneic setting (**Fig. 5j**). Some of the genes enriched in early rejection included S1pr5, linked to T-cell trafficking (*32*); Erdr1, associated with immune regulation (*52*); Pla2g4b, involved in inflammatory lipid metabolism and immune cell activity (*53*); and Clec2g, related to NK cell receptors (*54*), together suggesting heightened immune activation in the early rejection group. Principal component analysis of this signature demonstrated clear separation between the two outcome groups (**Fig. 5k**), indicating that IN captures early molecular programs linked to the timing of graft failure.

We generated an early-rejection score using gene set variation analysis based on relative enrichment of early rejection associated IN signature genes to track rejection dynamics over time (**Sup. Table 8)**. In early-rejection group, the score progressively increased and was already elevated by day 28, indicating imminent graft loss. In contrast, late-rejection group had an initial decline in the score through day 28, followed by a subsequent rise prior to graft failure (**Fig. 5l**). Temporal divergence and later re-emergence of this score highlight the ability of IN not only to distinguish rejection phenotypes, but also to longitudinally detect evolving immune risk before overt loss of graft function. We next examined pathway-level dynamics to investigate the biology underlying evolution of immune programs in early- and late-rejection groups. Notably, inflammation, TNF-NF-κB, interferon, IL-6-JAK-STAT, NK/NKT-cell, and T-cell signaling gene sets were higher at baseline in early-rejection group than in the late-rejection group, suggesting a more primed immune state before transplantation (**Fig. 5m**). Following transplantation and immunotherapy and rapamycin treatment, these pathways became comparable between groups at days 7 and 14, consistent with an initial therapeutic suppression of immune activation. However, by day 28, several pathways, including IL-6-JAK-STAT, NK/NKT, and T-cell signaling, showed an increase in early-rejection group, whereas rise in late-rejection group was delayed until closer to day 70. These temporal differences indicate that the early rejection group has faster increasing inflammatory and cytotoxic programs as compared to the late rejection group.

We next investigated the potential for IN to resolve immune components associated with alloimmune versus autoimmune responses to graft rejection under immunotherapy. Pathway enrichment profiles derived from rejection-associated IN genes in anti-CD40L treated allografts (rejection versus acceptance) were compared with those from anti-CD40L plus rapamycin treated auto-allografts (early versus late rejection) (**Fig. 5n, Sup. Table 4**). Interestingly, pathways related to TNF-α inflammation, inflammatory response, interferon signaling, CXCL9-associated macrophages, IL-6, neutrophils, and classical monocytes were similarly enriched in both allo-only and auto-allo settings, suggesting that these myeloid inflammatory programs are core features of alloimmune graft injury under immunotherapy. In contrast, pathways related to T cells, NK-cell cytotoxicity, B cells, and antigen presentation were preferentially enriched in auto-allo setting, indicating that added lymphoid component may reflect the autoimmune component of rejection. Together, these findings demonstrate that IN can deconvolute overlapping rejection biology and distinguish innate inflammatory programs primarily associated with alloimmunity from adaptive immune programs linked to autoimmunity.

## DISCUSSION

Monitoring of islet transplantation remains one of the most challenging areas in transplantation medicine. Unlike vascularized solid organs like heart or kidney that are transplanted as a discrete anatomical unit, islets are infused as dispersed cellular clusters, most commonly into hepatic portal circulation, where they engraft as microscopic deposits distributed throughout the liver (*55*). This diffuse and inaccessible architecture makes direct imaging and tissue biopsy highly challenging, preventing routine histological assessment of graft health (*17, 56*). Current clinical measures, including blood glucose, C-peptide, HbA1c, and glucose tolerance testing, act largely as functional readouts and often reflect damage after considerable loss of viable graft mass has already occurred (*57*). While approaches such as blood gene expression profiling and plasma donor-derived cell-free DNA have shown promise in other transplant settings, their application to islet transplantation remains constrained by low tissue mass, dispersed engraftment, and uncertain sensitivity for early injury (*58, 59*). Here, we report an engineered immunological niche that serves as a remote graft surrogate for minimally invasive longitudinal monitoring of transplant-associated immune responses in two clinically relevant models: allogeneic transplantation in C57BL/6 mice under anti-CD40L, and auto-alloimmune transplantation in NOD mice under anti-CD40L plus rapamycin.

The IN has strong potential to function as a surrogate tissue for transplanted islet grafts. The transcriptomic signature derived from IN that distinguished allogeneic from syngeneic transplantation was reflective of immune dysregulation. Several genes within this signature highlighted distinct graft environments. *Cd59b*, elevated in syngeneic transplantation, encodes a complement regulatory molecule that may protect islets from complement-mediated injury (*60*). *Malat1* and *Lncpint*, both linked to inflammatory responses, were also higher in syngeneic grafts, suggesting that even syngeneic transplantation induces an early inflammatory signaling (*61*). However, longitudinal analysis showed that this inflammation resolved over time in syngeneic group, suggesting tolerance. In contrast, the allogeneic group displayed a more adaptive immune-driven rejection profile. *S1pr5*, involved in lymphocyte trafficking, *Hlf*, associated with pro-inflammatory CD4+ T-cell activity, and *Ido2*, linked to pro-inflammatory B-cell responses, were enriched in allogeneic transplantation (*32–34*). Overall, IN gene signature-based scores were higher across most immune cell populations in allogeneic grafts than in syngeneic grafts, spanning both innate and adaptive compartments. These findings indicate that IN mirrors transcriptomics differences between allografts and syngeneic grafts and is indicative of graft-associated immune biology. Interestingly, IN also captured graft-associated immune cell and transcriptomic changes in graft acceptance vs rejection group in anti-CD40L treated allografts. IN identified macrophages and dendritic cells, particularly tissue-resident anti-inflammatory CX3CR1**+** subsets, as key immune populations related to allograft acceptance under anti-CD40L. These cells have established roles in pancreas islet homeostasis and transplant immunity, supporting biological relevance of the IN findings (*62, 63*). The acceptance-associated IN gene signature included *Pla2g2d, Cd5l*, and *Ifi30*, which are linked to immunoregulation, antigen handling, and M2-like macrophage polarization (*36, 39, 40*). Consistent with the IN signature, macrophages and dendritic cells were the only allograft cell types with significantly higher acceptance than rejection scores based on the IN-derived signature. IN also captured adaptive immune programs associated with autoimmune-mediated rejection in the auto-allograft setting, consistent with prior reports (*64*). Overall, these findings across multiple transplant settings support that IN captures tissue-associated immune changes reflective of transplanted islet graft. By capturing rejection-associated adaptive immune responses in untreated allografts, acceptance-associated regulatory myeloid programs under anti-CD40L therapy, and autoimmune rejection programs in the auto-allograft setting, the IN functions as a minimally invasive surrogate tissue for longitudinal monitoring of graft immune status and mechanisms underlying transplant outcomes.

IN provides comprehensive insights into immune programs that distinguish graft outcomes across transplantation settings, resolving signatures of rejection, tolerance, and therapeutic response under anti-CD40L immunotherapy. IN revealed a shift in immune landscape under anti-CD40L treatment, where rejection is less lymphoid-driven and more characterized by enrichment of innate immune populations, including neutrophils, macrophages, dendritic cells, and their associated subsets. This result is consistent with inhibition of T cell activation by anti-CD40L co-stimulatory blockade and aligns with prior studies highlighting the role of innate inflammation in islet transplantation (*65–67*). Macrophages and neutrophils have been shown to mediate allograft rejection through mechanisms such as phagocytosis and trogocytosis, while graft-infiltrating dendritic cells have promoted rejection by locally presenting donor alloantigens to effector T cells within the graft. Interestingly, in the anti-CD40L treated allograft acceptance group, IN identified subsets of these innate immune cells associated with regulatory and tolerogenic functions, suggesting a more nuanced role in graft outcomes depending on cellular subtypes. Previous reports have also highlighted the importance of regulatory dendritic cells and macrophages in promoting transplant tolerance under co-stimulatory blockade through immunoregulatory and T cell-suppressive mechanisms (*68, 69*). Together, these findings suggest that IN-derived immune signatures can resolve distinct innate immune subsets that differentially contribute to rejection versus tolerance. Importantly, IN also captures dynamic immune changes over time within acceptance group under anti-CD40L. Despite initial graft acceptance, a progressive increase in inflammatory innate immune signatures was observed, including neutrophil and macrophage-associated inflammation, relative to syngeneic controls. Notably, this increase is more evident by day 56 post-transplant in IN, whereas blood glucose levels do not rise until approximately day 100 in animals that eventually reject the graft. This time lag highlights the ability of IN to detect early, subclinical inflammatory changes that precede overt graft failure. Overall, IN distinguishes graft acceptance versus rejection while capturing anti-CD40L driven shifts in pro-inflammatory and regulatory innate immune programs, enabling prediction of transplant outcomes.

In conclusion, we demonstrate that synthetic IN acts as surrogate tissue for transplanted islet grafts, enabling minimally invasive monitoring of immune processes that determine islet graft fate in clinically relevant murine models. By identifying early rejection signatures, distinguishing underlying immune mechanisms, and capturing heterogeneity in autoimmune and allograft response to anti-CD40L immunotherapy, IN addresses major unmet diagnostic challenges in post-transplant care. Such an approach could allow and provide a window for intervention before irreversible graft loss and guide selection or adjustment of immunotherapies, while reducing reliance on highly invasive graft biopsies and delayed metabolic markers. Overall, these findings position IN as a new diagnostic paradigm in transplantation, where an accessible engineered tissue functions as an immune biopsy for early assessment of graft health to improve long-term patient outcomes.

## MATERIALS AND METHODS

### Animal Cohort Design

All animal studies were conducted in accordance with institutional guidelines and protocols approved by the University of Missouri Institutional Animal Care and Use Committee. For allogeneic islet transplant, BALB/c as donors and C57BL/6 recipients were used with or without anti-mouse CD40 ligand (CD40L) monoclonal antibody treatment. Syngeneic transplants utilized C57BL/6 mice for both donors and recipients. For auto-allogeneic studies, female non-obese diabetic (NOD/ShiLtJ) mice served as recipients of C57BL/6 donor islets, treated with a combination of anti-CD40L mAb and rapamycin. Individual mouse details are provided in **Sup. Table 1**.

### Immunological Niche Fabrication

Microporous poly(ε-caprolactone) (PCL) based immunological niches were fabricated by salt leaching technique as previously described (*70*). First 99g of NaCl (250 to 425um) was mixed with 1g of PCL pellets for 1hour at 85□°C. Subsequently, around 77.5 mg□of PCL-NaCl melt dispersion was pressed into 5mm scaffolds at 1500□psi for 45 seconds. They were then heated at 65 ° C for 5 mins on each side to anneal PCL. The scaffolds were placed in deionized water bath for salt leaching, and resulting porous niches were sanitized with 70% ethanol immersion followed by sterile PBS rinses.

### Immunological Niche implantation and retrieval

Mice were anesthetized with 2% isoflurane and maintained under anesthesia via a nose cone throughout all surgical procedures. INs were implanted into the subcutaneous space through a small dorsal incision. To minimize post-operative pain, mice received carprofen (5□mg/kg, subcutaneous) at time of surgery and again 24□hours post-operatively and were monitored during recovery for signs of distress. For IN retrieval, mice were re-anesthetized with 2% isoflurane, and the implant site was shaved and disinfected. A small incision adjacent to the implant was reopened using sterile forceps, and IN was carefully excised and removed. The incision was closed with wound clips. Explanted INs designated for transcriptomics analysis were immediately flash-frozen in isopentane, maintained on dry ice, and stored at -80□°C until further processing. INs intended for flow cytometry were immediately placed in MACS buffer (500mL 1X PBS, 8.3 mL of 30% BSA, 2 mL of 0.5M EDTA) and stored in ice prior to processing.

### Flow Cytometry

Explanted INs were processed into single-cell suspensions for flow cytometric analysis by mechanical dissociation followed by enzymatic digestion. Briefly, INs were minced into small fragments and digested in Liberase TL (Millipore Sigma, Cat. No. 5401020001; 10 U/mL) for 30 min at 37 °C. The digested material was passed through a 70-µm cell strainer, and red blood cells were lysed using ACK buffer (Fisher Scientific, Cat. No. A1049201) for 1 min. Cells were subsequently washed and resuspended in FACS buffer consisting of PBS supplemented with 0.5% bovine serum albumin and 2 mM EDTA. Cells were stained with multicolor antibody panels for immune phenotyping and acquired on a Cytek Northern Lights Aurora spectral flow cytometer. Flow cytometry data were analyzed using FlowJo v10.10.0. Samples collected at intermediate days were assigned to the nearest timepoint for analysis. Additional details regarding antibody panels, staining conditions, controls, and gating strategies are provided in the Supplementary Materials.

### Bulk RNA Sequencing Analysis

Frozen INs were homogenized in 800 µL TRIzol reagent (Fisher Scientific, Cat. No. 15596026) using mechanical disruption at 15,000 rpm. Total RNA was subsequently purified from the lysate using Direct-zol RNA Miniprep Plus kit (Fisher Scientific, Cat. No. NC1047980) according to manufacturer’s protocol. RNA quantity and purity were assessed prior to submission to the Advanced Genomics Core at the University of Michigan for bulk RNA sequencing. Sequencing libraries were prepared using poly-A enrichment workflows for most samples, with a subset processed using ribosomal RNA depletion-based library preparation. Libraries were sequenced on an Illumina NovaSeq X Plus platform. Raw base call files were converted to demultiplexed FASTQ files using BCL Convert software v4.0 (Illumina).

Bulk RNA-seq count matrices were quality filtered prior to downstream analysis. Samples collected at intermediate timepoints were assigned to the nearest analysis timepoint. Differential expression analysis was performed using DESeq2 with adjustment for batch, timepoint, and treatment effects where applicable (*71*). Variance-stabilized expression values were batch-corrected for downstream analyses while preserving group-level effects. Elastic net modeling with stability selection was used to derive transcriptomic signatures distinguishing experimental groups. Gene set enrichment analysis (GSEA) was performed using clusterProfiler with MSigDB C8 immune signatures, Hallmark and KEGG pathways (*72*). Gene set variation analysis (GSVA) was performed using curated immune-related gene sets on batch-corrected variance-stabilized expression data (*73, 74*). Additional details regarding preprocessing, elastic net optimization, stability selection, and score generation are provided in the supplementary methods.

### Single Cell RNA Sequencing Analysis

Published islet single-cell RNA-sequencing data from Chen et al. (*35*) were reanalyzed using Seurat v5.4.0 (*75*). Following quality control, normalization, clustering, and dimensionality reduction, IN-derived Elastic Net signatures were projected onto single-cell datasets using per-cell module scoring approaches. Single-sample gene set enrichment analysis (ssGSEA) was performed using MSigDB C8 gene sets, Hallmark and KEGG pathways. Additional details regarding single-cell preprocessing, module scoring, pathway analyses, and statistical testing are provided in the Supplementary Materials.

### Statistics

Statistical analyses for flow cytometry and blood glucose were performed using GraphPad Prism v10.6.1 and R. Diabetes-free survival was analyzed using Kaplan-Meier analysis. For two-group flow cytometry comparisons, unpaired t-tests or Mann-Whitney tests were used depending on normality. Longitudinal analyses were performed using mixed-effects models with multiple-comparison correction where applicable. Differential gene expression analysis was performed using DESeq2. For single-cell analyses, Wilcoxon rank-sum tests with Benjamini-Hochberg correction were used.

## Acknowledgments

Schematics were created in https://BioRender.com. Library prep and next-generation sequencing of INs was carried out in the Advanced Genomics Core at the University of Michigan.

## Funding

This work was supported by Breakthrough T1D (formerly JDRF) grants 2-SRA-2023-1454-S-B (L.D.S.) and 2-SRA-2024-1480-S-B (L.D.S). This work also received support from NIH grant U01EB036955.

## Author contributions

Conceptualization: JR, AJN, LDS, ESY

Methodology: JR, AJN, MT, AT, SW, BH, YJ, LDS, ESY

Investigation: JR, AJN, MT, AT, BH, LDS, ESY

Visualization: JR, AJN, LDS, ESY

Funding acquisition: LDS, ESY

Project administration: LDS, ESY

Supervision: LDS, ESY

Writing – original draft: JR, LDS, ESY

Writing – review & editing: JR, AJN, AT, BH, LDS, ESY

## Competing interests

Authors declare no competing interests.

## Data and materials availability

IN bulk sequencing data will be made available upon acceptance. Islet graft single cell RNA sequencing data can be found at Gene Expression Omnibus, accession number GSE198865. Codes used in this study is publicly available in Github: https://github.com/shea-lab/IsletTransplantRejection. All flow data will be available in the Deep Blue Repositories University of Michigan Library after acceptance.

## Notes

### Competing Interest Statement

The authors have declared no competing interest.

