## Supplementary Figures and Methods for "Immune Biomarkers of Islet Transplant Rejection Revealed by Synthetic Immunological Niche"

**List of Supplementary Materials**

**Supplementary Figures**

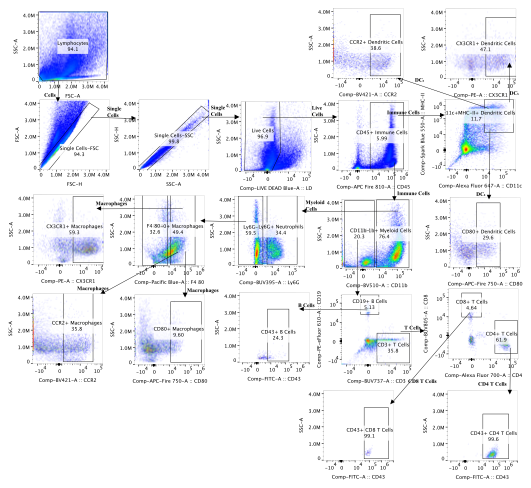

**Sup. Fig. 1: Representative Flow Gating Scheme:**  Gating strategy used for spectral flow cytometry analysis for the immunological niches from C57BL/6 transplants. Gates were slightly adjusted for NOD INs and liver tissues

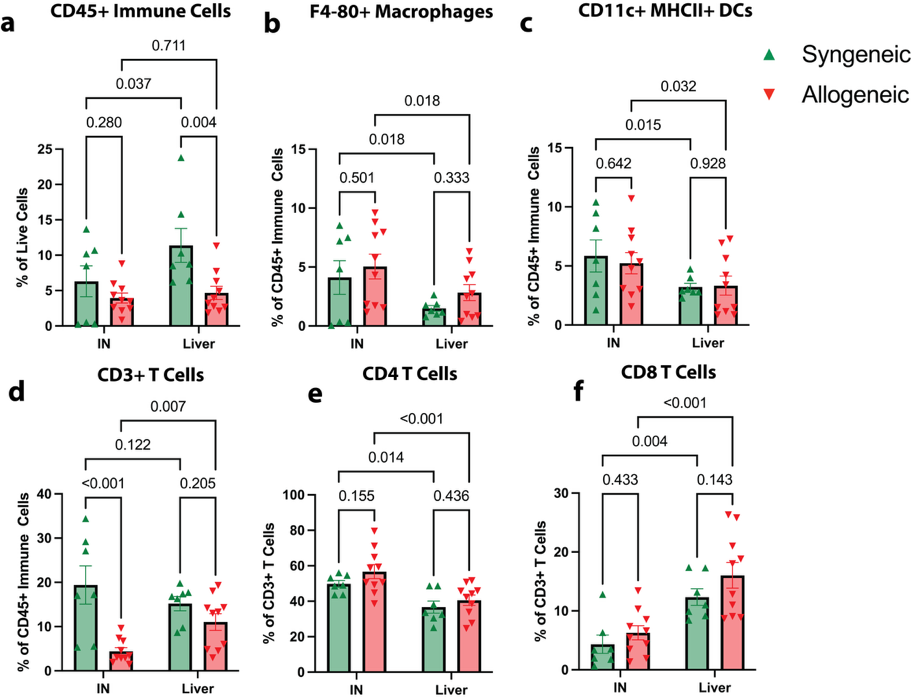

**Sup. Fig. 2: Immunophenotyping of IN and islet-transplanted liver at endpoint in allogeneic vs syngeneic groups.** Immune composition of the IN and islet-transplanted liver in syngeneic and allogeneic groups at their respective endpoints, including **(a)** total immune cells, **(b)** macrophages, **(c)** dendritic cells, **(d)** T cells, **(e)** CD4+ T cells, and **(f)** CD8+ T cells. These populations were detected in both IN and liver, with several cell types showing consistent trends between allogeneic and syngeneic conditions across the two sites. Statistical analysis was performed using a mixed-effects model with Fisher’s LSD (pooled variance).

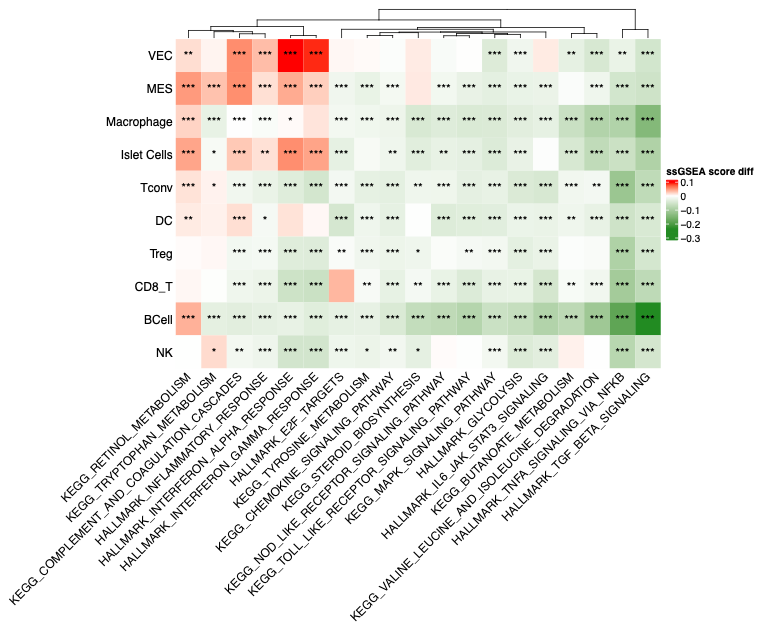

**Sup. Fig. 3: ssGSEA analysis of allogeneic vs syngeneic islet grafts.** Heatmap of differences in mean ssGSEA scores of selected IN-based genesets across cell types in allogeneic vs syngeneic islet grafts. Statistical significance was assessed using a Wilcoxon rank-sum test with Benjamini–Hochberg correction.

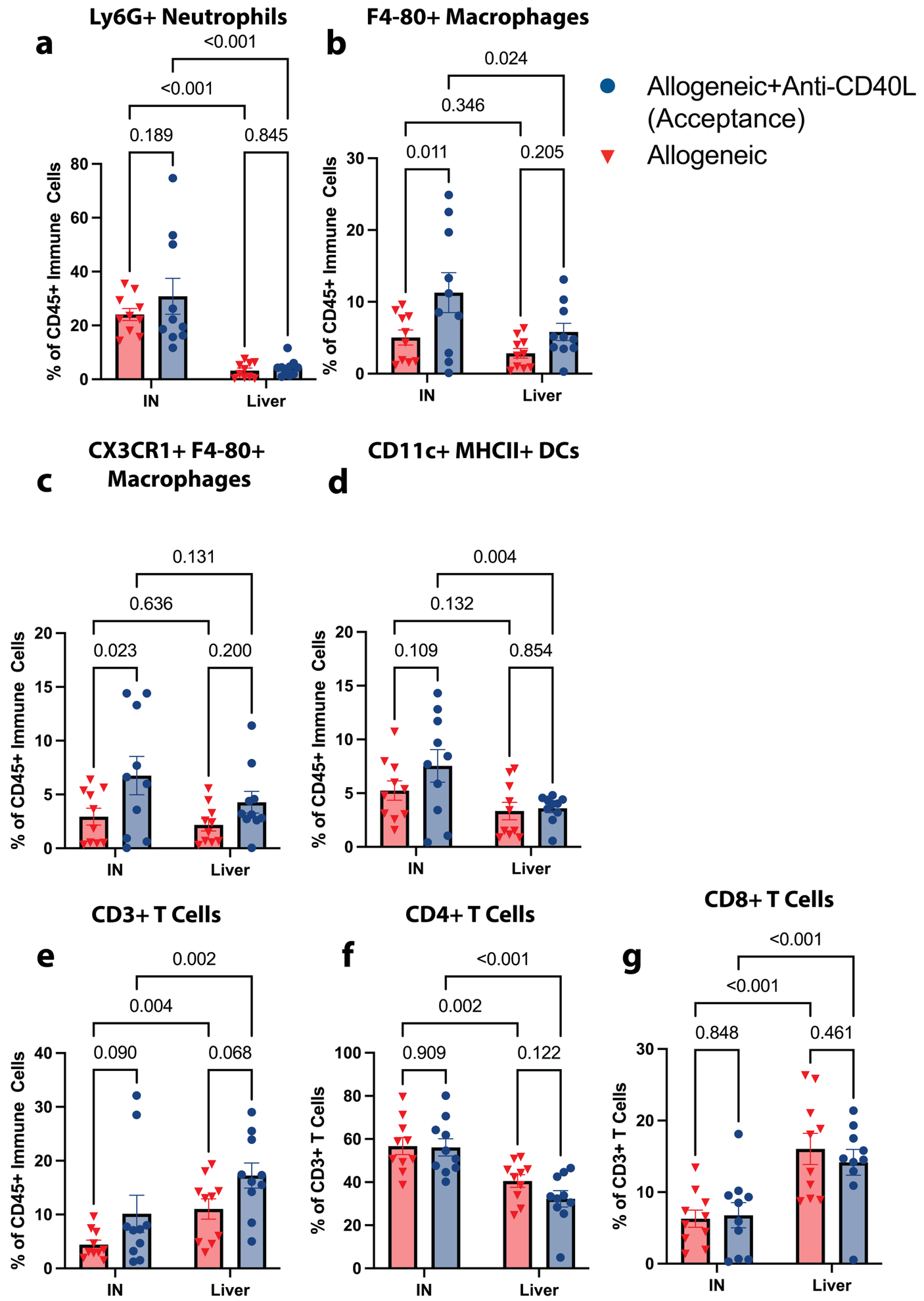

**Sup. Fig. 4: Immunophenotyping of IN and islet-transplanted liver at endpoint in allogeneic and allogeneic+anti-CD40L treated (accepted) groups.** Immune composition of the IN and islet-transplanted liver in allogeneic and allogeneic + anti-CD40L groups at their respective endpoint, including **(a)** neutrophils, **(b)** macrophages, **(c)** CX3CR1+ macrophages, **(d)** dendritic cells, **(e)** T cells, (**f)** CD4+ T cells, and **(g)** CD8+ T cells. These populations were detected in both IN and liver, with several cell types showing consistent trends between allogeneic and syngeneic conditions across the two tissues. Statistical analysis was performed using a mixed-effects model with Fisher’s LSD (pooled variance).

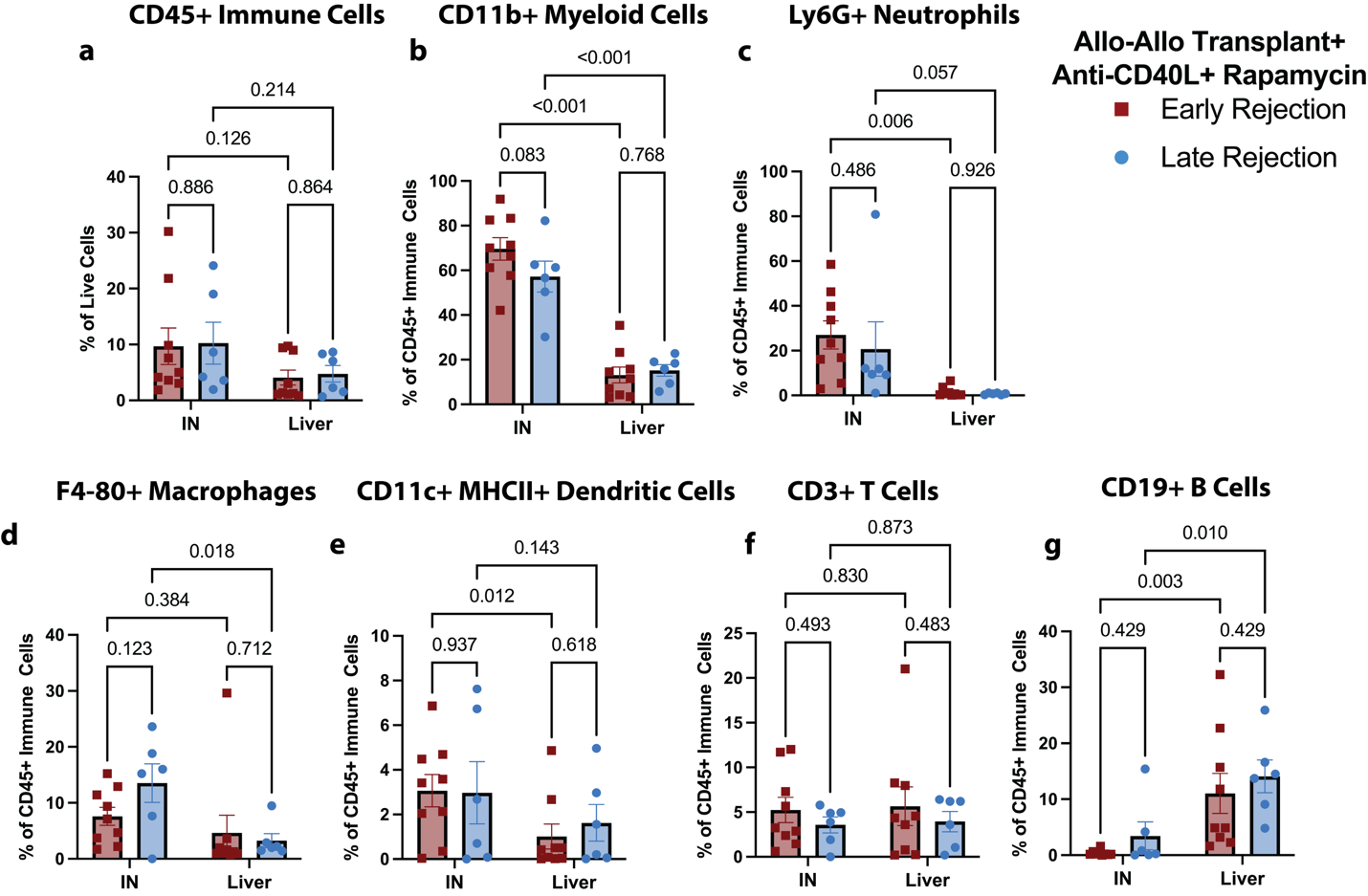

**Sup. Fig. 5: Immunophenotyping of IN and islet-transplanted liver at endpoint in early vs late rejection groups in NOD allogeneic islet transplants.** Immune composition of the IN and islet-transplanted liver in early vs late rejection groups in NOD allogeneic transplants treated with anti-CD40L and rapamycin at their respective endpoint, including **(a)** immune cells, **(b)** myeloid cells, **(c)** neutrophils, **(d)** macrophages, **(e)** dendritic cells, **(f)** T cells, and **(g)** B cells. These populations were detected in both IN and liver, with several cell types showing consistent trends between early vs late rejection groups across the two tissues. Statistical analysis was performed using a mixed-effects model with Fisher’s LSD (pooled variance).

**Supplementary methods**

**Flow Cytometry analysis**

For staining, cells were aliquoted at approximately 1 × 10^6 cells per tube. Samples were incubated with anti-CD16/32 Fc receptor blocking antibody (BD Bioscience, Cat. No. 553142; 1:50) for 15 min at 4 °C in dark, followed by surface staining for 30 min at 4 °C in dark with a multicolor antibody panel. Core markers were maintained across experiments, with fluorophore substitutions or panel refinements introduced as needed to optimize spectral resolution between runs. The panel included LIVE/DEAD Fixable Blue Dead Cell Stain (Thermo Fisher Scientific, Cat. No. L23105; 1:400), anti-mouse CD45 APC-Fire 810 (clone 30-F11, BioLegend, Cat. No. 103174; 1:150), anti-mouse CD3 (BUV737, clone 17A2, Thermo Fisher Scientific, Cat. No. 367-0032-82; 1:20 or V500 (clone 500A2, BD Horizon, Cat. No. 560771; 1:50) or APC, clone 145-2C11, BD Pharmingen, Cat. No. 553066, 1:50), anti-mouse CD4 Alexa Fluor 700 (clone RM4-5, Thermo Fisher Scientific, Cat. No. 56-0042-82; 1:150), anti-mouse CD19 PE-eFluor 610 (clone 1D3/CD19, Thermo Fisher Scientific, Cat. No. 61-0193-82; 1:100), anti-mouse F4/80 Pacific Blue (clone BM8, BioLegend, Cat. No. 123124; 1:100), anti-mouse CD11b BV510 (clone M1/70, BioLegend, Cat. No. 101263; 1:50), anti-mouse CD11c Alexa Fluor 647 (clone N418, BioLegend, Cat. No. 117312; 1:100), anti-mouse Ly6G BUV395 (clone 1A8, BD Biosciences, Cat. No. 563978; 1:100), anti-mouse CD80 APC-Fire 750 (clone 16-10A1, BioLegend, Cat. No. 104740; 1:50), anti-mouse I-A/I-E (MHC-II) Spark Blue 550 (clone M5/114.15.2, BioLegend, Cat. No. 107662; 1:50), anti-mouse CD43 FITC (clone S11, BioLegend, Cat. No. 143204; 1:20), anti-mouse Ly6C BV605 (clone HK1.4, BioLegend, Cat. No. 128036; 1:20), anti-mouse CX3CR1 PE (clone Z8-50, BD Biosciences, Cat. No. 567530; 1:50), anti-mouse CCR2 BV421 (clone SA203G11”, BioLegend, Cat. No. 150605; 1:100), and anti-mouse CD86 PECy7 (clone GL1, Thermo Fisher Scientific, Cat. No. 25-0862-82; 1:200). Following staining, cells were washed twice with PBS, resuspended in 300 µL PBS, and acquired on a Cytek Northern Lights Aurora flow cytometer. For each sample, 20,000 or more CD45+ events were collected. Flow cytometry data were analyzed using FlowJo v10.10.0. Single-stained and fluorescence-minus-one (FMO) controls were included for compensation and gating. Samples collected at intermediate days were assigned to the nearest timepoint for analysis.

**Bulk RNA sequencing analysis**

Bulk RNA-sequencing count matrices were preprocessed using a custom filtering pipeline for quality control and removal of low-confidence features, followed by exclusion of undefined transcripts and pseudogenes prior to downstream analysis. Samples collected at intermediate days were assigned to the nearest timepoint for analysis. Terminal samples collected after Day 56 were grouped with the Day 70 end-timepoint bin. Differential gene expression analysis was performed using DESeq2 with adjustment for batch, timepoint and treatment effects wherever applicable. For gene signature derivation, variance-stabilized gene expression values were batch-corrected while preserving group-level differences and adjusting for timepoint as a covariate. Candidate features were restricted to genes meeting differential expression thresholds (p ≤ 0.05 and |log_2_ FC| ≥ 0.5), reducing dimensionality prior to model fitting. An elastic net model was used to classify allogeneic versus syngeneic samples. The parameter α, which controls balance between L1 and L2 penalties in elastic net, was systematically evaluated across a predefined grid. For each α, regularization strength ($\lambda$) was optimized using leave-one-out cross-validation, with model performance assessed by cross-validated deviance. The optimal parameter $\alpha$ was selected based on minimizing deviance while maintaining enough retained features to balance predictive performance and model interpretability. To ensure robustness of feature selection, stability selection was performed over 1,000 iterations using repeated subsampling of approximately 80% of samples from each class to preserve class balance. Gene selection frequency was computed across iterations, and genes identified in ≥70% of runs were defined as stable features. For allogeneic versus syngeneic comparison, sex chromosome-associated genes were excluded from final signature. Gene set enrichment analysis (GSEA) was performed stage using clusterProfiler, with gene sets obtained from MSigDB, including Hallmark and KEGG pathway collections. All genes were ranked by the DESeq2 Wald statistic and pathways with adjusted p ≤ 0.1 and absolute normalized enrichment score (|NES|) ≥ 1 were considered significantly enriched. Gene set variation analysis (GSVA) was performed on variance-stabilized bulk RNA-seq data following batch correction using limma. For allogeneic acceptance vs syngeneic, GSVA scores were computed using a Gaussian kernel for curated and custom immune gene sets, including inflammatory macrophage, neutrophil, and T cell signatures. For the auto-allogeneic signature score, GSVA was applied separately to predefined upregulated and downregulated gene sets, and the composite score was calculated as difference between enrichment scores of upregulated and downregulated genes for each sample. Pathway enrichment scores were calculated per sample and compared across groups and time points using Welch’s t-test.

**Single cell RNA sequencing analysis**

Islet single-cell RNA-sequencing data from Chen et al. was reanalyzed using Seurat 5.4.0, with quality control filtering, SCTransform normalization, and PCA/UMAP-based dimensionality reduction. Clusters were identified using shared nearest neighbor graph-based clustering and annotated using previously defined canonical marker genes. To quantify IN-derived immune signatures differentiating allogeneic vs syngeneic in SCT-normalized single-cell data, an Elastic Net–derived gene signature was applied by computing a per-cell score as the difference between average expression of upregulated and downregulated genes in the IN signature. For the allogeneic graft single-cell data, rejection- and acceptance-associated Elastic Net gene signatures were scored separately using Seurat AddModuleScore, and module scores were compared across cell types. Single-sample gene set enrichment analysis (ssGSEA) was performed using curated Hallmark and KEGG pathways to estimate pathway activity at the single-cell level. Differences between conditions were quantified as mean pathway score differences and assessed using Wilcoxon rank-sum tests with Benjamini-Hochberg correction.
